# A whole-cell recording database of neuromodulatory action in the adult neocortex

**DOI:** 10.1101/2022.01.12.476007

**Authors:** Xuan Yan, Niccolo Calcini, Payam Safavi, Asli Ak, Koen Kole, Fleur Zeldenrust, Tansu Celikel

## Abstract

**Background:** The recent release of two large intracellular electrophysiological databases now allows high-dimensional systematic analysis of mechanisms of information processing in the neocortex. Here, to complement these efforts, we introduce a freely and publicly available database that provides a comparative insight into the role of various neuromodulatory transmitters in controlling neural information processing.

**Findings:** A database of *in vitro* whole-cell patch-clamp recordings from primary somatosensory and motor cortices (layers 2/3) of the adult mice (2-15 months old) from both sexes is introduced. A total of 464 current-clamp experiments from identified excitatory and inhibitory neurons are provided. Experiments include recordings with (i) Step-and-Hold protocol during which the current was transiently held at 10 steps, gradually increasing in amplitude, (ii) “Frozen Noise” injections that model the amplitude and time-varying nature of synaptic inputs to a neuron in biological networks. All experiments follow a within neuron across drug design which includes a vehicle control and a modulation of one of the following targets in the same neuron: dopamine and its receptors D1R, D2R, serotonin 5HT1f receptor, norepinephrine Alpha1, and acetylcholine M1 receptors.

**Conclusions:** This dataset is the first to provide a systematic and comparative insight into the role of the selected neuromodulators in controlling cellular excitability. The data will help to mechanistically address how bottom-up information processing can be modulated, providing a reference for studying neural coding characteristics and revealing the contribution of neuromodulation to information processing.

## Data description

Any behaviorally relevant computation in the brain is mediated through a network of neurons. Individual neurons, which constitute the nodes in the network, have powerful information processing capabilities ^1^. The process is governed by (i) neuronal properties, e.g. cell class, type and density of calcium buffers ^2,3^, (ii) biophysical properties of the cell, e.g. membrane capacitance, ion channel type, and density ^4,5^, as well as (iii) the nature of (chemical) communication impinging onto the neuron ^6,7^. Because neurons primarily integrate information sent to them through ligand-gated neurotransmission, understanding how different neuromodulators alter information processing is crucial to uncover the mechanism of information coding in the brain.

Even though Recent large-scale studies ^8,9^ now provide freely available datasets to biophysically and computationally study the mechanisms of information processing, they nonetheless do not address the contribution of different neurotransmitters, including neuromodulators, in this process. Neuromodulators, e.g. dopamine ^10^, serotonin ^11^, acetylcholine ^12^, and norepinephrine ^13^, as regulators of neuronal communication throughout the brain, control various neural, behavioural, and metabolic functions ^14,15^. The highly efficient and complex regulation of neuromodulators depends on their large family of receptor subtypes, receptor expression sites, ligands that are being synthesised and released, as well as the identity of the postsynaptic neurons ^7,16–18^. Here we provide a database collected from the neocortex of the mouse to help to contribute to these efforts. Focusing on the supragranular layers (Layers 2/3, L2/3) of the adult mouse, we intracellularly recorded from different types of neurons in sensory and motor cortices. Two protocols that mimic the sustained vs time-and-amplitude varying synaptic inputs were delivered to each identified neuron before and after pharmacological interventions targeting specific receptors. The Step-and-Hold protocol, i.e. a sustained depolarization of the membrane potential at 10 different depolarization steps, provides information regarding the active electrophysiological properties of the cell, including but not limited to the action potential threshold, spike timing, firing rate, and firing pattern, i.e. relative timing of action potentials in a spike train. The “frozen noise” protocol simulates the synaptic input impinging on a given postsynaptic cell, originating from 1000 synaptically connected presynaptic neurons ^1,3,19^ and allows for the quantification of intracellular information transfer as time-and-amplitude varying somatic currents are translated into action potential sequences. Both protocols are repeated multiple times while controlling the pharmacological state of the intervention in the neuron (see Materials and Methods).

The database is freely available online (<u>bit.ly/NeuromodulationDatabase</u>) and will help address: (1) the regulation of neuronal activity by neuromodulators through specific receptors, (2) neuromodulatory control of information processing and transfer under sustained and acute changes in neuronal excitability, (3) Cell-type specific control over intracellular information processing by neuromodulatory transmission. It will complement the recent efforts in the transcriptomic ^20,21^, proteomic ^22^, cellular ^8,23^, and behavioural ^1,24–26^ mapping of the sensory cortex to mechanistically study the neural information processing from molecules to behaviour.

## Materials and Method

### Experiments: Ethics statement

All experiments were conducted in accordance with the European Directive 2010/63/EU and have been approved by the local and national authorities prior to the start of experiments. Adult mice (Pvalbtm1(cre)Arb (RRID: MGI:5315557) or Ssttm2.1(cre)Zjh/J (RRID: IMSR_JAX:013044)) were backcrossed to C57BL6 and maintained on a 12-hour day-night cycle. They had ad libitum access to food and water and were kept in family cages until the day of the experiment.

### Experiments: Slicing preparations

All the procedures were as described before ^20,27–29^. Total of 232 cells were studied in vitro in coronal sections of the primary somatosensory and primary motor cortices of the mouse (Figure 1). On the day of the experiment, each animal was deeply anaesthetised with isoflurane before transcardially perfused using ice-cold physiological (slicing) solution containing (in mM) 108 choline chloride, 3 KCl, 26 NaHCO^3^, 1.25 NaH^2^PO^4^ ·H^2^O, 25 glucose·H^2^O, 1 CaCl^2^ ·2H^2^O, 6 MgSO^4^ ·7H^2^O. The brain was removed and placed in the same medium we used before. Coronal sections (300 micrometres) were cut from the barrel cortex subfield of the primary somatosensory cortex or the primary motor cortex while the brain was submerged in ice-cold slicing solution as described before ^20^. Sections were transferred to a recovery chamber filled with artificial cerebrospinal fluid (ACSF) containing (in mM) 120 NaCl, 3.5 KCl, 10 glucose·H^2^O, 2.5 CaCl^2^ ·2H^2^O, 1.3 MgSO^4^·7H^2^O, 25 NaHCO^3^, and 1.25 NaH^2^PO 4·H^2^O at 37 °C. After an hour of incubation, the slices were transferred to room temperature and used for recording. All the solutions used were pre-oxygenated with 95% O^2^ and 5% CO^2^.

**Figure 1.**
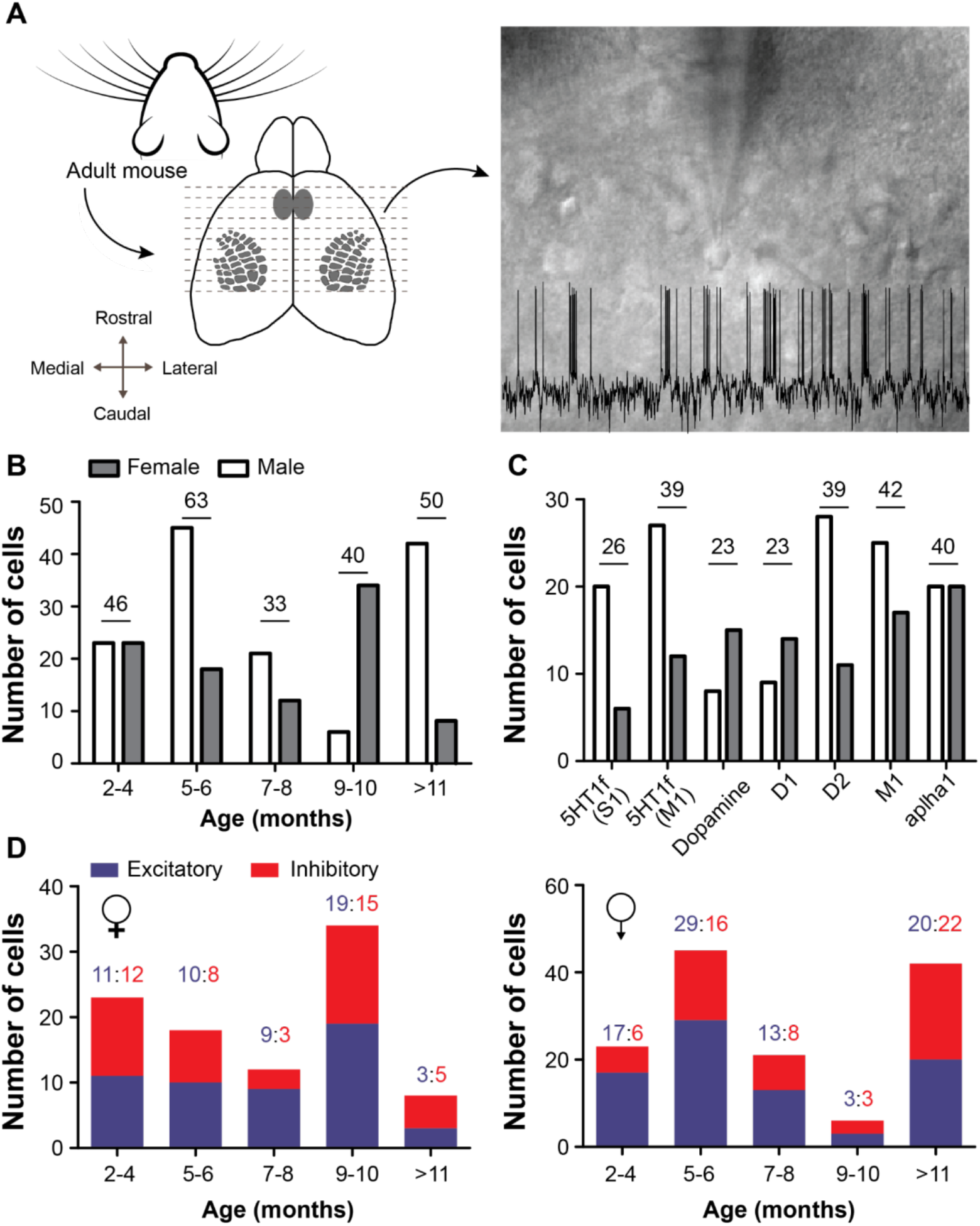
Description of the experimental procedures and dataset. Acute slices from the primary somatosensory (barrel) cortex and primary motor cortex, which is located rostra-laterally to the primary somatosensory cortex, were prepared from adult mice. Whole-cell current clamp recordings from putative excitatory and inhibitory neurons were made using two different stimulation conditions (Step- and-Hold and Frozen Noise, see Materials and Methods for details), before and during administration of receptor specific agonists (see Materials and Methods for details). **(A)** Left: Coronal slices from the adult neocortex were prepared. Right: Representative examples of microscopic imaging of soma at 40x and the membrane potential recorded (inset). **(B-D)** The distribution of the number of cells recorded across sexes (B), drug conditions and (C) cell classes.

### Experiments: Whole-cell recording

The slice was placed in a chamber filled with ACSF and was continuously oxygenated throughout recordings. The primary somatosensory or primary motor cortex was located under the 2.5X magnification and target neurons were visually identified at 40X. We use HEKA EPC10 amplifiers and Patch Master v2 × 90.2 software for recording. The glass capillaries (1.00 mm [external diameter], 0.50 mm [internal diameter], 75 mm [length], GC100FS-7.5, Harvard Apparatus) were used as the electrodes after they pulled using a P-2000 puller (Sutter Instrument, USA). The initial resistance was 5-9 MOhm. The intracellular solution contained (in mM) 130 K-Gluconate, 5 KCl, 1.5 MgCl^2^·6H^2^O, 0.4 Na^3^GTP, 4 Na^2^ATP, 10 HEPES, 10 Naphosphocreatine, and 0.6 EGTA, whose pH was set at 7.22 with KOH and was filled into the electrodes for patching.

Step-and-Hold protocol: we performed current-clamp recordings before and after the receptor activation to determine the basic biophysical and spiking properties of each neuron. The resting membrane potential was set at −70mV. Neurons whose resting membrane potential fluctuated >7mV throughout the experiment were excluded from the database. Current injections were delivered in 10 incremental steps of 20 or 40pA/step for 500 ms with an inter-sweep interval of 6.5s (see Figure 2).

**Figure 2.**
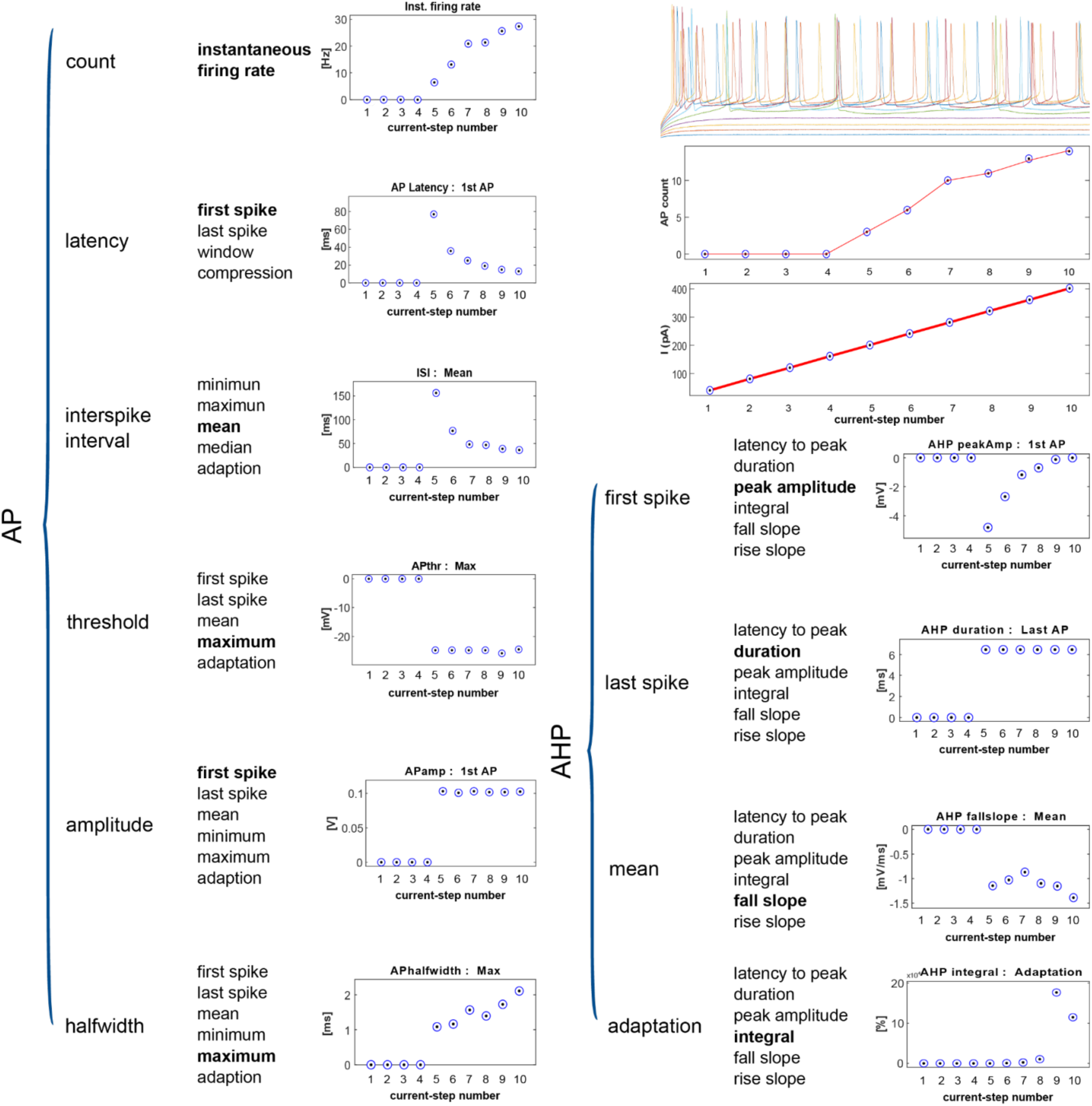
The representative dataset collected with the Step-and-Hold protocol. The variables are provided as a hierarchical tree. Variables in bold are the data displays shown on the right side of each column. AP=action potential, AHP=afterhyperpolarization. “Latency compression” is the difference in delay between the first and last action potential during each stimulation period normalised to the stimulus duration. “Adaptation” is the relative change of the observed variable normalised to the first event. The membrane potential traces on the top-right show the spiking responses to different amplitudes of injected current, superimposed on each other. The data underneath represents the number of action potentials and the intensity of current injected under 10 step-and-hold stimuli (Filename=asli_2-7-19-E2-CCSTEP-NODRUG; neuron class: excitatory).

Frozen Noise protocol: We established an artificial neural network composed of 1000 neurons, each firing Poisson spike trains in response to a ‘hidden state’, a binary signal switching between an ‘on’ and an ‘off’ state according to a Markov process^19^. For each recorded inhibitory or excitatory neuron, the same ‘frozen noise’ signal was used, although the switching speed of the hidden state and the firing rates of the artificial neurons differed across cell types (see Table 1). The fluctuating frozen noise current (amplitude: 700, 2100 pA, for excitatory and inhibitory neurons respectively) was injected in addition to a baseline current that kept resting membrane potential at −70mV, both before and after receptor activation. The resulting membrane potential and currents were recorded.

**Table 1.** Frozen noise parameters

| Parameter | Excitatory cells | Inhibitory cells |
| --- | --- | --- |
| Number of artificial neurons N | 1000 | 1000 |
| Hidden state time constant $\tau$ | 250 ms | 50 ms |
| Average firing rate artificial neurons $\mu_q$ | 0.1 Hz | 0.5 Hz |

### Pharmacology

We regulated the activation of serotonin 5HT1f receptors (LY334370: 5μM), norepinephrine Alpha1 (Cirazoline hydrochloride: 40μM), acetylcholine M1 (McA-N-343: 20μM), as well as dopamine D1 (SKF38393: 1μM) and D2 receptors (Quinpirole: 10μM) independently and simultaneously using dopamine (50μM). All chemicals were obtained from Sigma-Aldrich.

All experiments included were within cell controls. Recordings in ACSF served as vehicle control, as all drugs were dissolved in ACSF. Both recording protocols (i.e. Step-and-Hold and Frozen Noise) were repeated before and after drug administration. The recordings upon drug administration started 3 min after each drug was released into the circulating bath solution. After each experiment, the recording chamber was cleaned and the circulating bath solutions were kept in separate containers to ensure that there is no cross-drug contamination.

### Data organisation

The metadata contains information about the recorded animals, including the age, sex, strain, the date of the recording, the experiment number, and the target receptor (see Supplemental Table1). The experiments are named as prefix_date-experiment number-CCSTEP(or FN)-condition. All cells recorded from the same animal share the same experimental date. The current clamp data (CurrentClamp folder) contains two subfolders, “Step-and-Hold protocol” and “Frozen Noise protocol”. Each cell has two files, they have the same name although the suffix depends on the stimulus condition. Similar to the “Step-and-Hold protocol”, the “FrozenNoise” subfolder contains all the recordings with the data recorded under the frozen noise protocol.

Data collected using the “Step-and-Hold protocol” includes the injected current, the recorded voltage, and the timestamps for each variable. The recordings are named similarly using a template, i.e “Trace_a_b_c_d” where (a) is the cell number, (b) sequence of recording, (c) sweep number, and (d) the channel, i.e 1 for current injected, 2 is for voltage observed.

The “Frozen Noise protocol” subfolder is organised similarly to the other one. Each datafile contains the cell response (i.e voltage recordings upon injected current), hidden state ^19^ (binary stimulus), the input current trace, and a Matlab structural variable named “settings”. These ‘settings’ include the following information: condition (ACSF or the specific agonist), experimenter, baseline current (in pA) that is injected to keep the resting membrane potential at −70 mV, amplitude_scaling ^19^ (to control spiking rate), tau (the time constant of the average switching speed of the hidden state), mean_firing_rate (the mean firing rate of the artificial neurons, in kHz), sampling_rate (the acquisition rate, in kHz), duration (in ms) and FLAG_convert_to_amphere (a binary value that is 1 if the output was converted into ampere) ^8^.

## Cell type identification

We putatively classified neurons into excitatory and inhibitory subclasses based on the shape of soma and the pattern of spiking. The soma of excitatory neurons in acute slices appear more triangular and are larger than inhibitory neurons. This criterion is initially used to target neurons whose cell class was confirmed post-hoc using the spike patterns. The end-user can classify the neurons based on the active properties of the spiking to identify cellular classes of interest.

## Re-use potential

The database provided herein contains sub- and suprathreshold voltage dynamics of adult cortical neurons upon sustained or time and amplitude modulated somatic depolarization with and without the activation of selected neuromodulatory receptors. It complements the dataset previously made available from our group^8^, which focused on the electrical characterization of sensory neocortical neurons in the adult brain as well as the community efforts to make electrophysiological data freely available ^9,30^. The stimulus protocols provided in this dataset will help address fundamental questions like how the neuromodulators change action potential dynamics, shape and patterning (including adaptation), what are the cell type specific consequences of neuromodulation on cellular excitability, and what are the contribution of neuromodulators to intracellular information transfer and communication in synaptically coupled networks.

Spike pattern adaptation to sustained current injection is often used to classify neuron types ^8^. Excitatory neurons usually show a lower firing frequency than inhibitory neurons, rapidly adapting to prolonged stimulation. The data collected with the Step-and-Hold protocol provides means to systematically address how neuromodulators alter firing rate, relative and absolute action potential timing in a cell type specific manner (compare Figure 2 to Figure 3 for a representative case from excitatory and inhibitory neurons, respectively).

**Figure 3.**
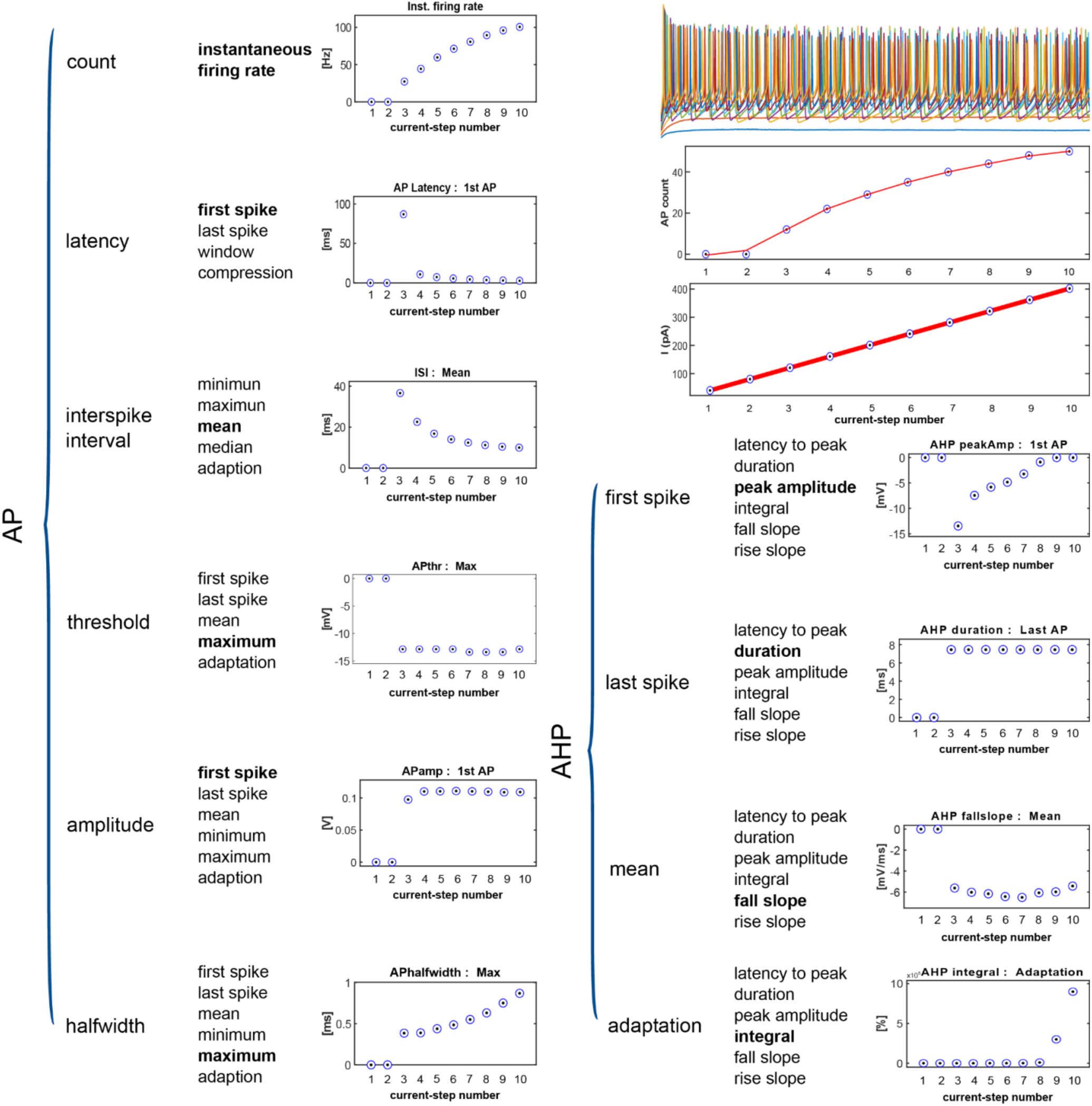
Responses of a representative inhibitory neuron to sustained current injection. Stimulus protocol: Step-and-Hold. Variables quantified are shown as a hierarchical tree. Those that are in bold are used as data displays on the right columns. Labels and descriptions of definitions are as in Figure 2. (Filename=asli_12-7-19-E3-CCSTEP-NODRUG).

The way neurons encode and process information, among other things, depends on the features of action potentials and spiking patterns ^31,32^ which reflect the activities of ion channels. While somatic depolarization is primarily regulated by calcium and sodium channels ^33^, repolarization is mediated by potassium and chloride channels ^34^. Neuromodulatory neurotransmitters could alter this process by phosphorylation of ion channels, thus controlling spiking patterns. The data provided herein will help address which neuromodulatory transmitters control action potential shape and its temporal dynamics, thus nominating receptor specific signalling pathways to explore the biophysical mechanism of neural information processing.

Spike patterns are the embodiment of information processing in the nervous system which is regulated by varying the rate and timing of action potentials ^1^. Quantification of intracellular information transfer from subthreshold to suprathreshold responses with and without the activation of neuromodulatory neurotransmitters will help shed light on the role of neuromodulation in information processing. The Frozen Noise protocol provides a systematic analysis of this process as time and amplitude varying somatic input is injected into the neuron while observing spike generation^8^. Researchers can analyse the changes between the input (current injected) and output (membrane potential recorded) to study information processing quantitatively. This approach ensures bias-free quantification of intracellular information transfer with a short (around 6 min) stimulation protocol ^19^.

Given the bandwidth limited time and amplitude varying nature of the current injection in the Frozen Noise protocol, the observations will complement the observations made with the help of the Step-and-Hold protocol, as researchers can directly compare the action potential dynamics during sustained and fluctuating depolarization with and without receptor intervention.

## Application prospects

This database will contribute to multidisciplinary efforts, serving biologists, physiologists, computational and systems neuroscientists. It advances global efforts for the systematic analysis of neural responses across species ^8,35,36^. Because our experimental protocols are identical across the database provided herein and the one we previously published ^8^, the user can benefit from a large dataset of intracellular recordings with over 1500 experiments performed in the adult brain. Developing a unified description of information processing across different types of neurons and identifying the common mechanisms that regulate information transfer across (different classes of) neurons will be a significant contribution to many branches of neuroscience.

The data provided herein can contribute to molecular physiology pipelines. Users can identify co-regulation networks in the transcriptome ^21,23,37^ and proteome ^22,38^ of the somatosensory cortex that we provided freely to the community; the data can be visualised online at barrelomics.science.ru.nl. By molecular analysis of transcription and translation for the neuromodulatory receptors regulated herein, the user can identify mechanisms of action that are coupled to action potential dynamics controlled by neuromodulation, and can study key genes and proteins that affect the coding and transmission of neuronal information.

The aforementioned molecular datasets include additional groups with experience dependent changes in the transcriptome and proteome after the altered sensory experience which ultimately control action potential dynamics as shown before ^25,27–29,39,40^. Differential transcriptome analysis (between sensory deprived and control conditions) for the downstream targets of the receptors modulated herein will help to identify the neuromodulatory mechanisms that contribute to adaptive changes in information processing.

Quantification of information transfer is traditionally performed using methods borrowed from signal processing literature. These methods are developed with different constraints than the experimental realities of intracellular recordings *in vitro*. Because neural recording quality is limited in duration, e.g. due to fluctuations in the membrane potential, drift, and change in the health of the cell in acute slices, electrophysiological recordings from individual neurons result in relatively short (<60 min) recording epochs that allow a limited number of synaptic input patterns to be studied in a given neuron. The Frozen Noise dataset provided in the current database as well as the one we made available previously ^8^ offer a solution.

## Limitations

The neurons in this database mainly come from neurons in the L2/3 layer of the sensory and motor cortices, thus the sample does not represent the spiking dynamics of neurons across all layers in distinct nuclei of the neocortex. We target neurons “blindly”, without any knowledge regarding their molecular cell type. Expanding these experiments with molecularly targeted cell populations will improve further classification of the neurons.

Due to the within cell, repeated measure design we cannot randomise the order of vehicle and drug conditions. Thus, in all experiments, vehicle control was delivered first. To control for this order bias, we have also provided experiments where the vehicle was delivered sequentially multiple times.

The analyses of both protocols (data not shown) show that the order bias does not impact the quality of recordings and the neural responses. Because we used each slice only once and the set-up was thoroughly cleaned in between recordings, there was no cross-drug contamination.

## Declarations

The authors declare that they have no competing interests.

## Funding

This work was supported by a doctoral fellowship from the Chinese Scholarship Council (CSC) to X.Y.; grants from the European Commission (Horizon2020, nr. 660328), European Regional Development Fund (MIND, nr. 122035), and the Netherlands Organisation for Scientific Research (NWO-ALW Open Competition, nr. 824.14.022) to T.C.; and by the Netherlands Organisation for Scientific Research (NWO Veni Research Grant, nr. 863.150.25) to F.Z.

**Supplemental Table1:** Metadata.

| *date*<br>(DD.MM.<br>YY) | *Experiment*<br>*number* | *Animal*<br>*number* | *prefix* | *Age(p)* | *sex* | *Experiment*<br>*protocol* | *Classification* | *Agonist* |
| --- | --- | --- | --- | --- | --- | --- | --- | --- |
| 30719 | E1 | A1 | xuan | 241 | M | Step-and-Hold,<br>Frozen Noise | Excitatory | 5HT-1f(S1<br>cortex) |
| 30719 | E2 | A1 | xuan | 241 | M | Step-and-Hold,<br>Frozen Noise | Excitatory | 5HT-1f(S1<br>cortex) |
| 30719 | E3 | A1 | xuan | 241 | M | Step-and-Hold,<br>Frozen Noise | Excitatory | 5HT-1f(S1<br>cortex) |
| 30719 | E4 | A1 | xuan | 241 | M | Step-and-Hold,<br>Frozen Noise | Excitatory | 5HT-1f(S1<br>cortex) |
| 70619 | E1 | A2 | xuan | 229 | F | Step-and-Hold,<br>Frozen Noise | Excitatory | 5HT-1f(S1<br>cortex) |
| 70619 | E2 | A2 | xuan | 229 | F | Step-and-Hold,<br>Frozen Noise | Excitatory | 5HT-1f(S1<br>cortex) |
| 70619 | E5 | A2 | xuan | 229 | F | Step-and-Hold,<br>Frozen Noise | Excitatory | 5HT-1f(S1<br>cortex) |
| 120619 | E1 | A3 | xuan | 145 | M | Step-and-Hold,<br>Frozen Noise | Excitatory | 5HT-1f(S1<br>cortex) |
| 120619 | E3 | A3 | xuan | 145 | M | Step-and-Hold,<br>Frozen Noise | Excitatory | 5HT-1f(S1<br>cortex) |
| 130619 | E4 | A4 | xuan | 146 | M | Step-and-Hold,<br>Frozen Noise | Excitatory | 5HT-1f(S1<br>cortex) |
| 140619 | E2 | A5 | xuan | 147 | M | Step-and-Hold,<br>Frozen Noise | Excitatory | 5HT-1f(S1<br>cortex) |
| 140619 | E3 | A5 | xuan | 147 | M | Step-and-Hold,<br>Frozen Noise | Excitatory | 5HT-1f(S1<br>cortex) |
| 140619 | E4 | A5 | xuan | 147 | M | Step-and-Hold,<br>Frozen Noise | Excitatory | 5HT-1f(S1<br>cortex) |
| 50619 | E4 | A6 | xuan | 227 | F | Step-and-Hold,<br>Frozen Noise | Excitatory | 5HT-1f(S1<br>cortex) |
| 260619 | E4 | A7 | xuan | 201 | M | Step-and-Hold,<br>Frozen Noise | Excitatory | 5HT-1f(S1<br>cortex) |
| 50619 | E3 | A6 | xuan | 227 | F | Step-and-Hold,<br>Frozen Noise | Inhibitory | 5HT-1f(S1<br>cortex) |
| 70619 | E4 | A2 | xuan | 229 | F | Step-and-Hold,<br>Frozen Noise | Inhibitory | 5HT-1f(S1<br>cortex) |
| 90719 | E3 | A8 | xuan | 215 | M | Step-and-Hold,<br>Frozen Noise | Inhibitory | 5HT-1f(S1<br>cortex) |
| 90719 | E4 | A8 | xuan | 215 | M | Step-and-Hold,<br>Frozen Noise | Inhibitory | 5HT-1f(S1<br>cortex) |
| 130619 | E1 | A4 | xuan | 146 | M | Step-and-Hold,<br>Frozen Noise | Inhibitory | 5HT-1f(S1<br>cortex) |
| 130619 | E2 | A4 | xuan | 146 | M | Step-and-Hold,<br>Frozen Noise | Inhibitory | 5HT-1f(S1<br>cortex) |
| 130619 | E3 | A4 | xuan | 146 | M | Step-and-Hold,<br>Frozen Noise | Inhibitory | 5HT-1f(S1<br>cortex) |
| 240619 | E1 | A9 | xuan | 199 | M | Step-and-Hold,<br>Frozen Noise | Inhibitory | 5HT-1f(S1<br>cortex) |
| 260619 | E2 | A7 | xuan | 201 | M | Step-and-Hold,<br>Frozen Noise | Inhibitory | 5HT-1f(S1<br>cortex) |
| 260619 | E6 | A7 | xuan | 201 | M | Step-and-Hold,<br>Frozen Noise | Inhibitory | 5HT-1f(S1<br>cortex) |
| 260619 | E5 | A7 | xuan | 201 | M | Step-and-Hold,<br>Frozen Noise | Inhibitory | 5HT-1f(S1<br>cortex) |
| 30419 | E2 | A10 | xuan | 236 | F | Step-and-Hold,<br>Frozen Noise | Excitatory | D2 |
| 30419 | E3 | A10 | xuan | 236 | F | Step-and-Hold,<br>Frozen Noise | Excitatory | D2 |
| 60319 | E1 | A11 | xuan | 230 | M | Step-and-Hold,<br>Frozen Noise | Excitatory | D2 |
| 60319 | E2 | A11 | xuan | 230 | M | Step-and-Hold,<br>Frozen Noise | Excitatory | D2 |
| 70319 | E2 | A12 | xuan | 231 | M | Step-and-Hold,<br>Frozen Noise | Excitatory | D2 |
| 70319 | E3 | A12 | xuan | 231 | M | Step-and-Hold,<br>Frozen Noise | Excitatory | D2 |
| 80119 | E2 | A13 | xuan | 47 | M | Step-and-Hold,<br>Frozen Noise | Excitatory | D2 |
| 80119 | E3 | A13 | xuan | 47 | M | Step-and-Hold,<br>Frozen Noise | Excitatory | D2 |
| 90119 | E3 | A14 | xuan | 48 | M | Step-and-Hold,<br>Frozen Noise | Excitatory | D2 |
| 101018 | E2 | A15 | xuan | 115 | F | Step-and-Hold,<br>Frozen Noise | Excitatory | D2 |
| 121018 | E1 | A16 | xuan | 117 | F | Step-and-Hold,<br>Frozen Noise | Excitatory | D2 |
| 121018 | E2 | A16 | xuan | 117 | F | Step-and-Hold,<br>Frozen Noise | Excitatory | D2 |
| 140319 | E2 | A17 | xuan | 238 | M | Step-and-Hold,<br>Frozen Noise | Excitatory | D2 |
| 151018 | E2 | A18 | xuan | 162 | M | Step-and-Hold,<br>Frozen Noise | Excitatory | D2 |
| 170119 | E2 | A19 | xuan | 140 | M | Step-and-Hold,<br>Frozen Noise | Excitatory | D2 |
| 170119 | E3 | A19 | xuan | 140 | M | Step-and-Hold,<br>Frozen Noise | Excitatory | D2 |
| 190319 | E4 | A20 | xuan | 211 | M | Step-and-Hold,<br>Frozen Noise | Excitatory | D2 |
| 200319 | E3 | A21 | xuan | 213 | F | Step-and-Hold,<br>Frozen Noise | Excitatory | D2 |
| 210119 | E1 | A22 | xuan | 152 | M | Step-and-Hold,<br>Frozen Noise | Excitatory | D2 |
| 210319 | E3 | A23 | xuan | 211 | M | Step-and-Hold,<br>Frozen Noise | Excitatory | D2 |
| 210319 | E4 | A23 | xuan | 211 | M | Step-and-Hold,<br>Frozen Noise | Excitatory | D2 |
| 221018 | E1 | A24 | xuan | 92 | F | Step-and-Hold,<br>Frozen Noise | Excitatory | D2 |
| 231018 | E1 | A25 | xuan | 93 | F | Step-and-Hold,<br>Frozen Noise | Excitatory | D2 |
| 231018 | E2 | A25 | xuan | 93 | F | Step-and-Hold,<br>Frozen Noise | Excitatory | D2 |
| 231118 | E2 | A26 | xuan | 128 | M | Step-and-Hold,<br>Frozen Noise | Excitatory | D2 |
| 240119 | E2 | A27 | xuan | 147 | M | Step-and-Hold,<br>Frozen Noise | Excitatory | D2 |
| 260319 | E2 | A28 | xuan | 185 | M | Step-and-Hold,<br>Frozen Noise | Excitatory | D2 |
| 260319 | E4 | A28 | xuan | 185 | M | Step-and-Hold,<br>Frozen Noise | Excitatory | D2 |
| 270319 | E2 | A29 | xuan | 186 | M | Step-and-Hold,<br>Frozen Noise | Excitatory | D2 |
| 280319 | E3 | A30 | xuan | 187 | M | Step-and-Hold,<br>Frozen Noise | Excitatory | D2 |
| 280319 | E4 | A30 | xuan | 187 | M | Step-and-Hold,<br>Frozen Noise | Excitatory | D2 |
| 290319 | E1 | A31 | xuan | 188 | M | Step-and-Hold,<br>Frozen Noise | Excitatory | D2 |
| 210119 | E5 | A22 | xuan | 152 | M | Step-and-Hold,<br>Frozen Noise | Inhibitory | D2 |
| 220119 | E1 | A32 | xuan | 153 | M | Step-and-Hold,<br>Frozen Noise | Inhibitory | D2 |
| 220119 | E2 | A32 | xuan | 153 | M | Step-and-Hold,<br>Frozen Noise | Inhibitory | D2 |
| 230119 | E1 | A33 | xuan | 146 | F | Step-and-Hold,<br>Frozen Noise | Inhibitory | D2 |
| 190319 | E1 | A20 | xuan | 211 | M | Step-and-Hold,<br>Frozen Noise | Inhibitory | D2 |
| 220319 | E2 | A34 | xuan | 212 | M | Step-and-Hold,<br>Frozen Noise | Inhibitory | D2 |
| 290319 | E2 | A31 | xuan | 188 | F | Step-and-Hold,<br>Frozen Noise | Inhibitory | D2 |
| 20719 | E2 | A35 | asli | 105 | M | Step-and-Hold,<br>Frozen Noise | Excitatory | M1 |
| 20819 | E1 | A36 | asli | 162 | M | Step-and-Hold,<br>Frozen Noise | Excitatory | M1 |
| 50719 | E3 | A37 | asli | 173 | M | Step-and-Hold,<br>Frozen Noise | Excitatory | M1 |
| 50719 | E4 | A37 | asli | 173 | M | Step-and-Hold,<br>Frozen Noise | Excitatory | M1 |
| 50819 | E1 | A38 | asli | 163 | F | Step-and-Hold,<br>Frozen Noise | Excitatory | M1 |
| 50819 | E2 | A38 | asli | 163 | F | Step-and-Hold,<br>Frozen Noise | Excitatory | M1 |
| 50819 | E3 | A38 | asli | 163 | F | Step-and-Hold,<br>Frozen Noise | Excitatory | M1 |
| 50819 | E7 | A38 | asli | 163 | F | Step-and-Hold,<br>Frozen Noise | Excitatory | M1 |
| 60819 | E2 | A39 | asli | 164 | F | Step-and-Hold,<br>Frozen Noise | Excitatory | M1 |
| 60819 | E5 | A39 | asli | 164 | F | Step-and-Hold,<br>Frozen Noise | Excitatory | M1 |
| 80719 | E2 | A40 | asli | 176 | M | Step-and-Hold,<br>Frozen Noise | Excitatory | M1 |
| 120719 | E2 | A41 | asli | 178 | M | Step-and-Hold,<br>Frozen Noise | Excitatory | M1 |
| 170619 | E2 | A42 | asli | 113 | F | Step-and-Hold,<br>Frozen Noise | Excitatory | M1 |
| 230719 | E2 | A44 | asli | 136 | M | Step-and-Hold,<br>Frozen Noise | Excitatory | M1 |
| 230719 | E3 | A44 | asli | 136 | M | Step-and-Hold,<br>Frozen Noise | Excitatory | M1 |
| 230719 | E5 | A44 | asli | 136 | M | Step-and-Hold,<br>Frozen Noise | Excitatory | M1 |
| 240719 | E5 | A45 | asli | 137 | M | Step-and-Hold,<br>Frozen Noise | Excitatory | M1 |
| 270619 | E1 | A46 | asli | 132 | M | Step-and-Hold,<br>Frozen Noise | Excitatory | M1 |
| 270619 | E2 | A46 | asli | 132 | M | Step-and-Hold,<br>Frozen Noise | Excitatory | M1 |
| 300719 | E1 | A47 | asli | 141 | M | Step-and-Hold,<br>Frozen Noise | Excitatory | M1 |
| 160719 | E1 | A48 | xuan | 144 | M | Step-and-Hold,<br>Frozen Noise | Excitatory | M1 |
| 160719 | E5 | A48 | xuan | 144 | M | Step-and-Hold,<br>Frozen Noise | Excitatory | M1 |
| 20819 | E2 | A36 | asli | 162 | M | Step-and-Hold,<br>Frozen Noise | Inhibitory | M1 |
| 20819 | E3 | A36 | asli | 162 | M | Step-and-Hold,<br>Frozen Noise | Inhibitory | M1 |
| 50819 | E4 | A38 | asli | 163 | F | Step-and-Hold,<br>Frozen Noise | Inhibitory | M1 |
| 60819 | E1 | A39 | asli | 164 | F | Step-and-Hold,<br>Frozen Noise | Inhibitory | M1 |
| 60819 | E3 | A39 | asli | 164 | F | Step-and-Hold,<br>Frozen Noise | Inhibitory | M1 |
| 60819 | E4 | A39 | asli | 164 | F | Step-and-Hold,<br>Frozen Noise | Inhibitory | M1 |
| 120719 | E3 | A41 | asli | 178 | M | Step-and-Hold,<br>Frozen Noise | Inhibitory | M1 |
| 170619 | E1 | A42 | asli | 113 | F | Step-and-Hold,<br>Frozen Noise | Inhibitory | M1 |
| 180719 | E1 | A43 | asli | 180 | F | Step-and-Hold,<br>Frozen Noise | Inhibitory | M1 |
| 240719 | E4 | A45 | asli | 137 | M | Step-and-Hold,<br>Frozen Noise | Inhibitory | M1 |
| 250719 | E1 | A49 | asli | 138 | M | Step-and-Hold,<br>Frozen Noise | Inhibitory | M1 |
| 250719 | E2 | A49 | asli | 138 | M | Step-and-Hold,<br>Frozen Noise | Inhibitory | M1 |
| 250719 | E3 | A49 | asli | 138 | M | Step-and-Hold,<br>Frozen Noise | Inhibitory | M1 |
| 250719 | E4 | A49 | asli | 138 | M | Step-and-Hold,<br>Frozen Noise | Inhibitory | M1 |
| 310719 | E1 | A50 | asli | 119 | F | Step-and-Hold,<br>Frozen Noise | Inhibitory | M1 |
| 310719 | E2 | A50 | asli | 119 | F | Step-and-Hold,<br>Frozen Noise | Inhibitory | M1 |
| 310719 | E3 | A50 | asli | 119 | F | Step-and-Hold,<br>Frozen Noise | Inhibitory | M1 |
| 310719 | E4 | A50 | asli | 119 | F | Step-and-Hold,<br>Frozen Noise | Inhibitory | M1 |
| 160719 | E2 | A48 | xuan | 144 | M | Step-and-Hold,<br>Frozen Noise | Inhibitory | M1 |
| 160719 | E3 | A48 | xuan | 144 | M | Step-and-Hold,<br>Frozen Noise | Inhibitory | M1 |
| 120919 | E4 | A60 | payam | 148 | F | Step-and-Hold,<br>Frozen Noise | Excitatory | Alpha1 |
| 260919 | E1 | A61 | payam | 113 | M | Step-and-Hold,<br>Frozen Noise | Excitatory | Alpha1 |
| 260919 | E2 | A61 | payam | 113 | M | Step-and-Hold,<br>Frozen Noise | Excitatory | Alpha1 |
| 21019 | E2 | A62 | payam | 90 | M | Step-and-Hold,<br>Frozen Noise | Excitatory | Alpha1 |
| 21019 | E3 | A62 | payam | 90 | M | Step-and-Hold,<br>Frozen Noise | Excitatory | Alpha1 |
| 31019 | E1 | A63 | payam | 91 | M | Step-and-Hold,<br>Frozen Noise | Excitatory | Alpha1 |
| 31019 | E3 | A63 | payam | 91 | M | Step-and-Hold,<br>Frozen Noise | Excitatory | Alpha1 |
| 31019 | E4 | A63 | payam | 91 | M | Step-and-Hold,<br>Frozen Noise | Excitatory | Alpha1 |
| 31019 | E5 | A63 | payam | 91 | M | Step-and-Hold,<br>Frozen Noise | Excitatory | Alpha1 |
| 101019 | E3 | A64 | payam | 128 | F | Step-and-Hold,<br>Frozen Noise | Excitatory | Alpha1 |
| 161019 | E4 | A65 | payam | 115 | M | Step-and-Hold,<br>Frozen Noise | Excitatory | Alpha1 |
| 120919 | E1 | A60 | payam | 148 | F | Step-and-Hold,<br>Frozen Noise | Excitatory | Alpha1 |
| 90919 | E2 | A67 | xuan | 366 | M | Step-and-Hold,<br>Frozen Noise | Excitatory | Alpha1 |
| 160919 | E2 | A68 | xuan | 376 | F | Step-and-Hold,<br>Frozen Noise | Excitatory | Alpha1 |
| 200919 | E2 | A69 | xuan | 377 | F | Step-and-Hold,<br>Frozen Noise | Excitatory | Alpha1 |
| 200919 | E3 | A69 | xuan | 377 | F | Step-and-Hold,<br>Frozen Noise | Excitatory | Alpha1 |
| 270919 | E1 | A70 | xuan | 271 | F | Step-and-Hold,<br>Frozen Noise | Excitatory | Alpha1 |
| 111019 | E3 | A71 | xuan | 211 | F | Step-and-Hold,<br>Frozen Noise | Excitatory | Alpha1 |
| 120919 | E3 | A60 | payam | 148 | F | Step-and-Hold,<br>Frozen Noise | Inhibitory | Alpha1 |
| 50919 | E1 | A72 | payam | 363 | M | Step-and-Hold,<br>Frozen Noise | Inhibitory | Alpha1 |
| 50919 | E2 | A72 | payam | 363 | M | Step-and-Hold,<br>Frozen Noise | Inhibitory | Alpha1 |
| 50919 | E3 | A72 | payam | 363 | M | Step-and-Hold,<br>Frozen Noise | Inhibitory | Alpha1 |
| 240919 | E1 | A73 | payam | 348 | F | Step-and-Hold,<br>Frozen Noise | Inhibitory | Alpha1 |
| 250919 | E1 | A74 | payam | 349 | F | Step-and-Hold,<br>Frozen Noise | Inhibitory | Alpha1 |
| 250919 | E2 | A74 | payam | 349 | F | Step-and-Hold,<br>Frozen Noise | Inhibitory | Alpha1 |
| 21019 | E1 | A62 | payam | 90 | M | Step-and-Hold,<br>Frozen Noise | Inhibitory | Alpha1 |
| 21019 | E6 | A62 | payam | 90 | M | Step-and-Hold,<br>Frozen Noise | Inhibitory | Alpha1 |
| 101019 | E2 | A64 | payam | 128 | F | Step-and-Hold,<br>Frozen Noise | Inhibitory | Alpha1 |
| 101019 | E4 | A64 | payam | 128 | F | Step-and-Hold,<br>Frozen Noise | Inhibitory | Alpha1 |
| 101019 | E5 | A64 | payam | 128 | F | Step-and-Hold,<br>Frozen Noise | Inhibitory | Alpha1 |
| 161019 | E3 | A65 | payam | 115 | M | Step-and-Hold,<br>Frozen Noise | Inhibitory | Alpha1 |
| 161019 | E5 | A65 | payam | 115 | M | Step-and-Hold,<br>Frozen Noise | Inhibitory | Alpha1 |
| 90919 | E1 | A67 | xuan | 366 | M | Step-and-Hold,<br>Frozen Noise | Inhibitory | Alpha1 |
| 90919 | E3 | A67 | xuan | 366 | M | Step-and-Hold,<br>Frozen Noise | Inhibitory | Alpha1 |
| 90919 | E4 | A67 | xuan | 366 | M | Step-and-Hold,<br>Frozen Noise | Inhibitory | Alpha1 |
| 160919 | E4 | A68 | xuan | 376 | F | Step-and-Hold,<br>Frozen Noise | Inhibitory | Alpha1 |
| 200919 | E1 | A69 | xuan | 377 | F | Step-and-Hold,<br>Frozen Noise | Inhibitory | Alpha1 |
| 270919 | E2 | A70 | xuan | 271 | F | Step-and-Hold,<br>Frozen Noise | Inhibitory | Alpha1 |
| 270919 | E3 | A70 | xuan | 272 | F | Step-and-Hold,<br>Frozen Noise | Inhibitory | Alpha1 |
| 111019 | E1 | A71 | xuan | 211 | F | Step-and-Hold,<br>Frozen Noise | Inhibitory | Alpha1 |
| 260520 | E2 | A75 | xuan | 358 | M | Step-and-Hold,<br>Frozen Noise | Excitatory | 5HT1f(M1<br>cortex) |
| 260520 | E3 | A75 | xuan | 358 | M | Step-and-Hold,<br>Frozen Noise | Excitatory | 5HT1f(M1<br>cortex) |
| 270520 | E3 | A76 | xuan | 359 | M | Step-and-Hold,<br>Frozen Noise | Excitatory | 5HT1f(M1<br>cortex) |
| 280520 | E2 | A77 | xuan | 340 | M | Step-and-Hold,<br>Frozen Noise | Excitatory | 5HT1f(M1<br>cortex) |
| 10620 | E3 | A78 | xuan | 291 | F | Step-and-Hold,<br>Frozen Noise | Excitatory | 5HT1f(M1<br>cortex) |
| 20620 | E1 | A79 | xuan | 292 | F | Step-and-Hold,<br>Frozen Noise | Excitatory | 5HT1f(M1<br>cortex) |
| 20620 | E5 | A79 | xuan | 292 | F | Step-and-Hold,<br>Frozen Noise | Excitatory | 5HT1f(M1<br>cortex) |
| 30620 | E1 | A80 | xuan | 293 | F | Step-and-Hold,<br>Frozen Noise | Excitatory | 5HT1f(M1<br>cortex) |
| 30620 | E2 | A80 | xuan | 293 | F | Step-and-Hold,<br>Frozen Noise | Excitatory | 5HT1f(M1<br>cortex) |
| 30620 | E3 | A80 | xuan | 293 | F | Step-and-Hold,<br>Frozen Noise | Excitatory | 5HT1f(M1<br>cortex) |
| 30620 | E4 | A80 | xuan | 293 | F | Step-and-Hold,<br>Frozen Noise | Excitatory | 5HT1f(M1<br>cortex) |
| 90620 | E2 | A81 | xuan | 313 | F | Step-and-Hold,<br>Frozen Noise | Excitatory | 5HT1f(M1<br>cortex) |
| 100620 | E4 | A82 | xuan | 314 | F | Step-and-Hold,<br>Frozen Noise | Excitatory | 5HT1f(M1<br>cortex) |
| 110620 | E3 | A83 | xuan | 324 | M | Step-and-Hold,<br>Frozen Noise | Excitatory | 5HT1f(M1<br>cortex) |
| 140620 | E1 | A84 | xuan | 327 | M | Step-and-Hold,<br>Frozen Noise | Excitatory | 5HT1f(M1<br>cortex) |
| 140620 | E3 | A84 | xuan | 327 | M | Step-and-Hold,<br>Frozen Noise | Excitatory | 5HT1f(M1<br>cortex) |
| 300620 | E1 | A85 | xuan | 343 | M | Step-and-Hold,<br>Frozen Noise | Excitatory | 5HT1f(M1<br>cortex) |
| 30720 | E2 | A86 | xuan | 364 | M | Step-and-Hold,<br>Frozen Noise | Excitatory | 5HT1f(M1<br>cortex) |
| 20720 | E3 | A87 | xuan | 363 | M | Step-and-Hold,<br>Frozen Noise | Excitatory | 5HT1f(M1<br>cortex) |
| 250520 | E2 | A88 | xuan | 357 | M | Step-and-Hold,<br>Frozen Noise | Inhibitory | 5HT1f(M1<br>cortex) |
| 260520 | E4 | A75 | xuan | 358 | M | Step-and-Hold,<br>Frozen Noise | Inhibitory | 5HT1f(M1<br>cortex) |
| 260520 | E5 | A75 | xuan | 358 | M | Step-and-Hold,<br>Frozen Noise | Inhibitory | 5HT1f(M1<br>cortex) |
| 280520 | E1 | A77 | xuan | 358 | M | Step-and-Hold,<br>Frozen Noise | Inhibitory | 5HT1f(M1<br>cortex) |
| 10620 | E4 | A78 | xuan | 291 | F | Step-and-Hold,<br>Frozen Noise | Inhibitory | 5HT1f(M1<br>cortex) |
| 10620 | E5 | A78 | xuan | 291 | F | Step-and-Hold,<br>Frozen Noise | Inhibitory | 5HT1f(M1<br>cortex) |
| 20620 | E3 | A79 | xuan | 292 | F | Step-and-Hold,<br>Frozen Noise | Inhibitory | 5HT1f(M1<br>cortex) |
| 40620 | E2 | A89 | xuan | 308 | M | Step-and-Hold,<br>Frozen Noise | Inhibitory | 5HT1f(M1<br>cortex) |
| 40620 | E5 | A89 | xuan | 308 | M | Step-and-Hold,<br>Frozen Noise | Inhibitory | 5HT1f(M1<br>cortex) |
| 50620 | E2 | A90 | xuan | 310 | M | Step-and-Hold,<br>Frozen Noise | Inhibitory | 5HT1f(M1<br>cortex) |
| 110620 | E1 | A83 | xuan | 324 | M | Step-and-Hold,<br>Frozen Noise | Inhibitory | 5HT1f(M1<br>cortex) |
| 110620 | E2 | A83 | xuan | 324 | M | Step-and-Hold,<br>Frozen Noise | Inhibitory | 5HT1f(M1<br>cortex) |
| 300620 | E4 | A85 | xuan | 343 | M | Step-and-Hold,<br>Frozen Noise | Inhibitory | 5HT1f(M1<br>cortex) |
| 10720 | E1 | A91 | xuan | 362 | M | Step-and-Hold,<br>Frozen Noise | Inhibitory | 5HT1f(M1<br>cortex) |
| 10720 | E2 | A91 | xuan | 362 | M | Step-and-Hold,<br>Frozen Noise | Inhibitory | 5HT1f(M1<br>cortex) |
| 10720 | E3 | A91 | xuan | 362 | M | Step-and-Hold,<br>Frozen Noise | Inhibitory | 5HT1f(M1<br>cortex) |
| 10720 | E4 | A91 | xuan | 362 | M | Step-and-Hold,<br>Frozen Noise | Inhibitory | 5HT1f(M1<br>cortex) |
| 20720 | E1 | A87 | xuan | 363 | M | Step-and-Hold,<br>Frozen Noise | Inhibitory | 5HT1f(M1<br>cortex) |
| 30720 | E1 | A86 | xuan | 364 | M | Step-and-Hold,<br>Frozen Noise | Inhibitory | 5HT1f(M1<br>cortex) |
| 30720 | E3 | A86 | xuan | 364 | M | Step-and-Hold,<br>Frozen Noise | Inhibitory | 5HT1f(M1<br>cortex) |
| 100519 | E1 | A51 | xuan | 259 | F | Step-and-Hold,<br>Frozen Noise | Excitatory | dopamine |
| 100519 | E2 | A51 | xuan | 259 | F | Step-and-Hold,<br>Frozen Noise | Excitatory | dopamine |
| 100519 | E3 | A51 | xuan | 259 | F | Step-and-Hold,<br>Frozen Noise | Excitatory | dopamine |
| 100519 | E4 | A51 | xuan | 259 | F | Step-and-Hold,<br>Frozen Noise | Excitatory | dopamine |
| 140519 | E2 | A52 | xuan | 339 | M | Step-and-Hold,<br>Frozen Noise | Excitatory | dopamine |
| 140519 | E4 | A52 | xuan | 339 | M | Step-and-Hold,<br>Frozen Noise | Excitatory | dopamine |
| 150519 | E2 | A54 | xuan | 340 | M | Step-and-Hold,<br>Frozen Noise | Excitatory | dopamine |
| 160519 | E3 | A55 | xuan | 307 | F | Step-and-Hold,<br>Frozen Noise | Excitatory | dopamine |
| 160519 | E5 | A55 | xuan | 307 | F | Step-and-Hold,<br>Frozen Noise | Excitatory | dopamine |
| 240519 | E1 | A56 | xuan | 312 | F | Step-and-Hold,<br>Frozen Noise | Excitatory | dopamine |
| 70519 | E1 | A57 | xuan | 281 | M | Step-and-Hold,<br>Frozen Noise | Excitatory | dopamine |
| 80519 | E4 | A58 | xuan | 282 | M | Step-and-Hold,<br>Frozen Noise | Excitatory | dopamine |
| 80519 | E5 | A58 | xuan | 282 | M | Step-and-Hold,<br>Frozen Noise | Excitatory | dopamine |
| 90519 | E4 | A59 | xuan | 258 | F | Step-and-Hold,<br>Frozen Noise | Excitatory | dopamine |
| 100519 | E5 | A51 | xuan | 258 | F | Step-and-Hold,<br>Frozen Noise | Inhibitory | dopamine |
| 100519 | E6 | A51 | xuan | 259 | F | Step-and-Hold,<br>Frozen Noise | Inhibitory | dopamine |
| 140519 | E1 | A52 | xuan | 339 | M | Step-and-Hold,<br>Frozen Noise | Inhibitory | dopamine |
| 150519 | E1 | A54 | xuan | 340 | M | Step-and-Hold,<br>Frozen Noise | Inhibitory | dopamine |
| 160519 | E1 | A55 | xuan | 307 | F | Step-and-Hold,<br>Frozen Noise | Inhibitory | dopamine |
| 160519 | E2 | A55 | xuan | 307 | F | Step-and-Hold,<br>Frozen Noise | Inhibitory | dopamine |
| 240519 | E4 | A56 | xuan | 312 | F | Step-and-Hold,<br>Frozen Noise | Inhibitory | dopamine |
| 90519 | E3 | A59 | xuan | 258 | F | Step-and-Hold,<br>Frozen Noise | Inhibitory | dopamine |
| 90519 | E1 | A59 | xuan | 258 | F | Step-and-Hold,<br>Frozen Noise | Inhibitory | dopamine |
| 110717 | E4 | A92 | NC | 89 | M | Step-and-Hold,<br>Frozen Noise | Excitatory | D1 |
| 120717 | E2 | A93 | NC | 84 | F | Step-and-Hold,<br>Frozen Noise | Excitatory | D1 |
| 150817 | E4 | A94 | NC | 118 | F | Step-and-Hold,<br>Frozen Noise | Excitatory | D1 |
| 210817 | E3 | A95 | NC | 84 | F | Step-and-Hold,<br>Frozen Noise | Excitatory | D1 |
| 300817 | E3 | A96 | NC | 285 | F | Step-and-Hold,<br>Frozen Noise | Excitatory | D1 |
| 101017 | E4 | A97 | NC | 335 | M | Step-and-Hold,<br>Frozen Noise | Excitatory | D1 |
| 101017 | E5 | A97 | NC | 335 | M | Step-and-Hold,<br>Frozen Noise | Excitatory | D1 |
| 111017 | E1 | A98 | NC | 372 | M | Step-and-Hold,<br>Frozen Noise | Excitatory | D1 |
| 111017 | E3 | A98 | NC | 372 | M | Step-and-Hold,<br>Frozen Noise | Excitatory | D1 |
| 111017 | E4 | A98 | NC | 372 | M | Step-and-Hold,<br>Frozen Noise | Excitatory | D1 |
| 121017 | E6 | A99 | NC | 373 | M | Step-and-Hold,<br>Frozen Noise | Excitatory | D1 |
| 171017 | E2 | A100 | NC | 154 | F | Step-and-Hold,<br>Frozen Noise | Excitatory | D1 |
| 171017 | E4 | A100 | NC | 154 | F | Step-and-Hold,<br>Frozen Noise | Excitatory | D1 |
| 201217 | E4 | A101 | NC | 218 | F | Step-and-Hold,<br>Frozen Noise | Excitatory | D1 |
| 300817 | E6 | A96 | KK | 285 | F | Step-and-Hold,<br>Frozen Noise | Inhibitory | D1 |
| 260617 | E1 | A102 | NC | 113 | M | Step-and-Hold,<br>Frozen Noise | Inhibitory | D1 |
| 260617 | E2 | A102 | NC | 113 | M | Step-and-Hold,<br>Frozen Noise | Inhibitory | D1 |
| 150817 | E1 | A94 | NC | 118 | F | Step-and-Hold,<br>Frozen Noise | Inhibitory | D1 |
| 150817 | E3 | A94 | NC | 118 | F | Step-and-Hold,<br>Frozen Noise | Inhibitory | D1 |
| 160817 | E1 | A103 | NC | 79 | F | Step-and-Hold,<br>Frozen Noise | Inhibitory | D1 |
| 160817 | E2 | A103 | NC | 79 | F | Step-and-Hold,<br>Frozen Noise | Inhibitory | D1 |
| 300817 | E1 | A96 | NC | 285 | F | Step-and-Hold,<br>Frozen Noise | Inhibitory | D1 |
| 140917 | E1 | A104 | NC | 309 | F | Step-and-Hold,<br>Frozen Noise | Inhibitory | D1 |

